# Hard-to-sample species are more sensitive to land-use change: implications for global biodiversity metrics

**DOI:** 10.1101/2025.01.16.633254

**Authors:** Claudia Gutiérrez-Arellano, Tim Newbold, Jenny A. Hodgson

## Abstract

Land-use change drives biodiversity loss, but some species suffer more than others. Indicators of global biodiversity change must attempt to summarise these impacts representatively and meaningfully, to guide biodiversity recovery. Yet species that are hard to detect, and thus feature less in relevant databases, might possess traits that make them particularly sensitive to anthropogenic impacts. Using global data for plant, bird, and spider species, we develop a statistical approach to analyse and correct for the impact of excluding hard-to-sample species from global biodiversity indicators. Based on over 4000 species with abundance comparisons available, we found that species with fewer global occurrence records consistently decline more as land-use intensity increases, suggesting that hard-to-sample species are particularly sensitive to land-use differences. When we extrapolate this relationship to all plant, bird and spider species with valid occurrence records (0.27 M species), we obtain a more representative global indicator of overall land-use impacts for these entire taxonomic groups. Our estimates indicate a lower average abundance in anthropogenic land uses compared to results obtained when hard-to-sample species are excluded. For example, intensive agriculture only has 18% of the biodiversity level of primary vegetation, rather than the 47% estimated without extrapolation. We recommend that other existing indicators include an extrapolation solution based on ours, to incorporate the available data as effectively as possible. Using occurrence data to predict species’ sensitivity unlocks many possibilities to improve global biodiversity indicators, without demanding additional data on poorly known species.

## Introduction

Land-use change is a major driver of biodiversity change, mainly through habitat loss and degradation (Purvis 2019; Jaureguiberry *et al*. 2022). A worldwide picture of how some aspects of biodiversity respond to land-use change seems to be within reach (Hill *et al*. 2018; Leclère *et al*. 2020; De Palma *et al*. 2021; Pereira *et al*. 2024), based on high-level indicators regarding suitable habitat extent, species richness, community composition, or relative abundance. Such indicators play an important role in the formulation and evaluation of conservation policies (Nicholson *et al*. 2019; Leclère *et al*. 2020; Ledger *et al*. 2023). However, species vary widely in their sensitivity to land-use change (Newbold *et al*. 2018; Sykes *et al*. 2020). To accurately represent global trends of change, models and indicators must strive to capture the wide range of responses among species and taxonomic groups (Jones *et al*. 2011; Hill *et al*. 2016).

In any one biodiversity metric, it is challenging to capture the aggregate effect across the ‘winners’ and ‘losers’. Of the influential global biodiversity metrics (as compared by Rosa *et al*. 2020) all use different tweaks either to reduce the influence of the abundant winners or to assume that the unimpacted state is the best one. This could be justified on the basis that the metrics are supposed to guide conservation, and conservation needs to focus on the subset of biodiversity that is threatened. However, it is possible tweaks might not be needed if the set of species analysed was truly representative. While biases in biodiversity data collection are well recognised (Beck *et al*. 2014; Di Marco *et al*. 2017), no one has developed a feasible way of correcting high-level indicators to account for the missing species.

Of course, some species are harder to sample than others; there are several species- and observer-related factors that reduce the likelihood of species turning up in biodiversity datasets (Hudson *et al*. 2014; Arazy & Malkinson 2021; Bennett *et al*. 2024). Missing species would not be a problem if their responses to anthropogenic threats were similar to those of the recorded species, but two factors contribute to a strong suspicion that this may not be the case. Firstly, evaluating the effect of different land uses requires standardised surveys of small plots. These surveys will tend to miss species that are cryptic or “rare” in different ways (Bennett *et al*. 2024) because of logistical constraints on survey effort, sampling methods and the sampled area. Secondly, looking within groups for which we already have a lot of data (i.e. vertebrate species), it seems that the rarer members are more likely to suffer from anthropogenic land uses (e.g. Newbold *et al*. 2018; Sykes *et al*. 2020). It is possible that traits that make species difficult to sample also make species more sensitive to land-use change. If so, by under-recording data from these highly sensitive species we will under-estimate biodiversity loss.

The lack of information for most species requires creative approaches to account for imperfect detection in biodiversity metrics and indicators (Bennett *et al*. 2024). Creative approaches may use the strengths of different data sources to complement one another (Twining *et al*. 2024). The strengths of occurrence records are their ubiquity and broad spatial coverage, providing data from the greatest possible range of species and landscapes. We propose the number of occurrence records as a metric for the ‘recordability’ of species, which includes all the factors known to lead to higher data volumes for some species than others. Number of records is likely to correlate to the chance of a species occurring in a wide range of ecological data sets, because of intuitive links to detectability (Arazy & Malkinson 2021) and abundance in some taxa (Callaghan *et al*. 2023). In parallel, its correlation with abundance and range size makes it a reasonable candidate as an indicator of sensitivity to land-use change (Newbold *et al*. 2018; Sykes *et al*. 2020).

To test the relationship between recordability and land-use sensitivity, we leveraged two global biodiversity databases, the PREDICTS (Projecting Responses of Ecological Diversity In Changing Terrestrial Systems, Hudson *et al*. 2016) and GBIF (Global Biodiversity Information Facility, https://www.gbif.org) databases. GBIF is the international collaborative database of species occurrence records in time and space. Although this database has taxonomic, geographic and temporal biases (Beck *et al*. 2014; Rocha-Ortega *et al*. 2021), it is currently the most comprehensive source of presence records globally — covering ∼1.75 M species (excluding unreviewed scientific names) of which at least 1.4 M of them have one occurrence record. The PREDICTS database (as updated in November 2022,Contu *et al*. 2022) collates over four million observations from studies that have compared the biodiversity of sites in different land use types and/or intensities. Its structure facilitates global analyses, and it is used to derive a number of indicators of the strength of anthropogenic impacts.

We explored the extent to which the relationship between site-level abundance and land-use type in PREDICTS is mediated by the number of records in GBIF for a species. We chose to include species from three taxonomic groups that cover a range of species richness and intensity of study: birds, plants and spiders. We found that species with fewer occurrence records are consistently more impacted by higher use intensities. Therefore, we were able to extrapolate this relationship to unstudied species, present in GBIF but not in PREDICTS (assuming that the relationships between recordability and the traits that directly relate to sensitivity hold for unstudied species). We hope that this highly adaptable approach can improve global biodiversity estimates while avoiding additional data collection.

## Materials and Methods

### PREDICTS database

We obtained the species’ local abundance data from the PREDICTS dataset, combining the data released in 2016 (Hudson *et al*. 2016) and 2022 (Contu *et al*. 2022). This joint dataset contains 4.3 million observations from 817 studies assessing the effects of land use change and intensification on approximately 32,000 species around the world. PREDICTS study sites are classified into nine predominant land uses and three levels of use intensity. Full definitions of predominant land use, the nine land use categories and use intensity can be found in the supplementary information of Hudson *et al*. (2014).

Briefly, land uses are:

- Primary vegetation: native vegetation that is not known or inferred to have ever been completely destroyed, before the year in which the biodiversity was sampled.
- Secondary vegetation: land where the original primary vegetation was completely destroyed. This is further divided into young, intermediate, mature or indeterminate depending on the structural complexity of the vegetation defined in the study.
- Plantation forest: land where people have planted crop trees or crop shrubs for commercial or subsistence harvesting.
- Pasture: land where livestock is known to be grazed regularly or permanently.
- Cropland: land where people have planted herbaceous crops, for human and livestock consumption.
- Urban: include areas with human habitation and/or buildings, where vegetation is managed for amenity purposes.

Use intensity is based on the level of disturbance — and extent of impact for primary and secondary vegetation— and is divided into light, minimal and intense. The exact definition varies depending on the land use type.

We simplified this land-use categorisation (Table S1) by following the classification suggested by Outhwaite *et al*. (2022), to obtain an easy-to-interpret gradient of increasing anthropogenic impact. This high-level classification of land use and use intensity (hereafter ‘land use’) includes primary vegetation, secondary vegetation, low-intensity agriculture and high-intensity agriculture. The studies included in our analyses were those that assessed two or more of these simplified land use types. We used the observations of those studies where the taxa were identified at the species level, the diversity metric type recorded was abundance, and there was a comparison of two or more of the simplified land-use types. This reduced the observation sample to 1.7 million records collated from 413 studies (Table S2).

We selected the records of three contrasting taxonomic groups: birds, a well-studied group with relatively low richness; spiders, a relatively unstudied species-rich group; and vascular plants, a relatively well-studied and species-rich group. Although these groupings are at different taxonomic levels (Class Aves for birds, order Araneae for spiders and subphylum Trachaeophyta for plants), they represent the level at which ecological surveys are most commonly organised. In some cases, included studies did sample some individuals outside the focal taxonomic group: in these cases, the non-relevant species were excluded from our analysis, but the rest of the study was kept.

### GBIF occurrence frequency

We obtained the number of records of all bird, plant and spider species in GBIF, referred to as occurrence count (Table S2). We filtered the data of those species for which their taxonomic status was accepted, were not extinct or extinct in the wild and their occurrence count was greater than zero before 2023 (period 1600-2023), which yielded 273, 420 species. The vast majority correspond to plant species (89%), 8% to spiders, and 3% to birds. Most of the species have less than 1000 records and a large proportion have less than 100 (Fig. 1a). In this sample, most of the number of records ranged from the thousands to the hundreds of thousands (min= 1; max = 22,679,448; mean= 156,399; median = 2,652; IQR =27,474; Fig. 1a). Before analysis, we log_10-_transformed the number of records and rescaled by subtracting the mean of this sample (Table S3).

**Figure 1.**
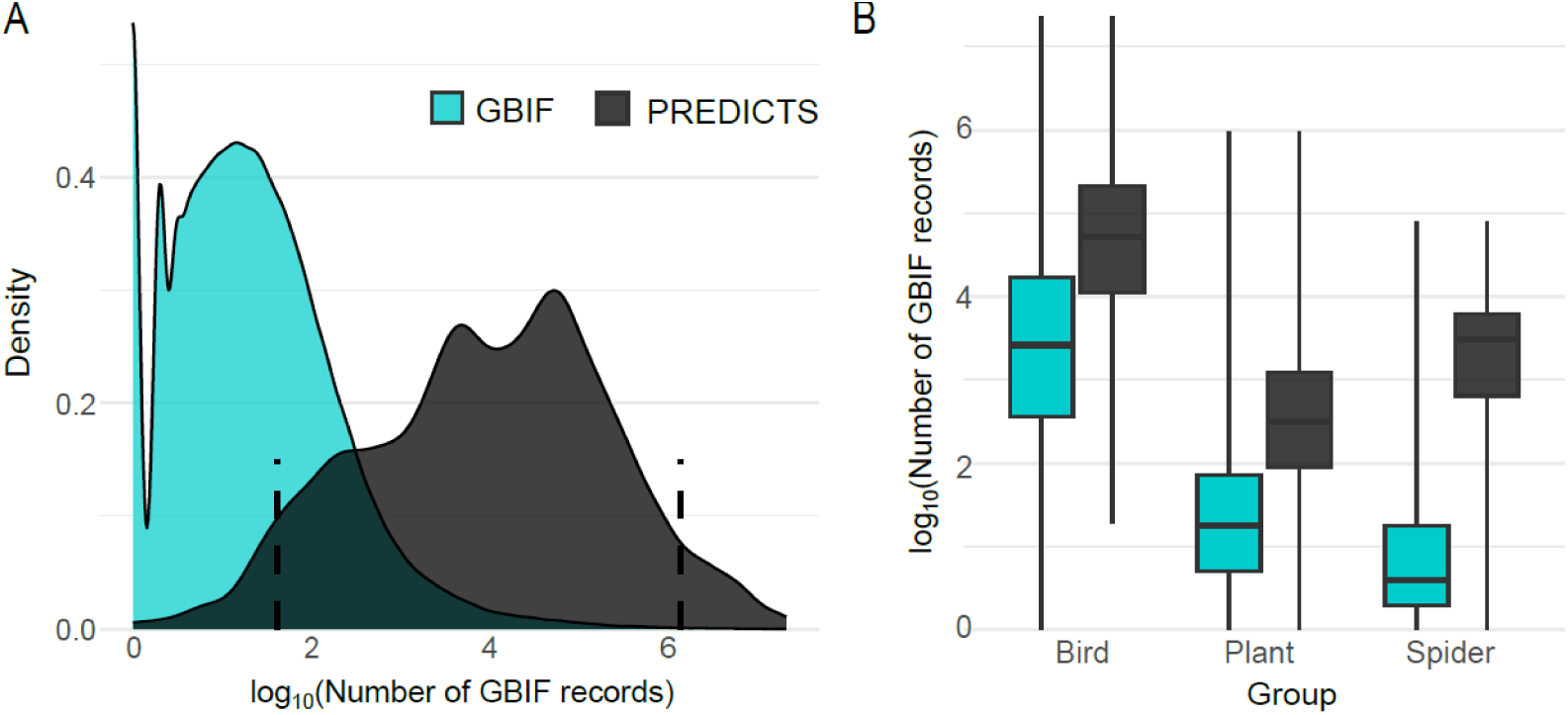
Distributions of the number of GBIF occurrence records across species. (A) Distributions as a density plot comparing all species available in GBIF (cyan, n=273,420 bird, plant, and spider species) to the species also present in our focal studies in the PREDICTS database (grey, n=4,454). Dashed lines indicate the 0.05 (42 records, left) and 0.95 (1,467,493 records, right) percentiles of the PREDICTS subset. (B) Species represented in GBIF and PREDICTS by taxonomic group of interest, with colours as in (A). The boxes show the quartiles, and the whiskers extend from minimum to maximum.

We matched the number of records to the species found in the PREDICTS database, based on PREDICTS’ Best Guess Binomial attribute (the inferred species’ scientific name, see Hudson *et al*. 2014), and found 4,454 matches (2,188 birds; 1,916 plants; 350 spiders; Table S2). Over a third of these species (1,568 spp.) were the subject of more than one study, the rest (65%) were present in one study. All three taxonomic groups of interest have all four land-use types represented (Fig. S1). The most common comparison within these studies was Primary vegetation vs. Secondary vegetation (28%) followed by Primary vegetation vs. Low-intensity agriculture (23%), while the least number of studies (12%) compared Secondary vegetation vs. High-intensity agriculture (Table S4).

### Total abundance and effort

For each PREDICTS study, we calculated the total abundance per species at each land use type (i.e. summing across any sites with the same simplified land use type)). This aggregation reduced the observations from 1.7 million to 67,000 observations of total abundance. Over half of these observations (35,637) belonged to bird, plant and spider species (Table S2). Total abundance values in our sample were right-skewed with a high number of zeros. Before analysis, we added the study’s minimum observed abundance to every abundance value and log_10-_transformed it. Similarly, we calculated the total survey effort per land cover type per study and log_10-_transformed it.

### Mixed effects model

We produced a linear mixed-effects model, hereafter the ‘Records model’, using the subset of the PREDICTS database records described above. We used the log_10-_transformed total abundance (total abundance) as the response variable, with land-use type (landuse), taxonomic group (taxon) and the transformed number of species records (records) as fixed effects (Table 1). Interaction terms allowed the effect of land use to depend on the number of records, and for the effects of records to vary between taxonomic groups (see model formula in Table 1). Study and species within study were random effects, and the log_10-_transformed total effort (total effort) per land-cover type per study was an offset term, meaning that the abundance of each species observed is assumed to be directly proportional to survey effort. We compared the goodness-of-fit of the Records model to alternative models — successively reducing the role played by the number of records (Table S5)— and the Records model was the best.

**Table 1.** Summary of ‘Records model’^1^ explaining the local abundance of species observed in PREDICTS. ‘landuse’ is an ordered factor, ordered from lowest to highest use intensity, fitted with polynomial contrasts. ‘taxon’ is an unordered factor and the transformed number of GBIF records ‘records’ is a scalar. Number of observations = 18,698; Random intercepts were fitted for studies (141 levels), and species within studies (7,763 levels). Note that because the local abundances for different taxa (and studies) tended to be measured in different units, this model’s coefficients can’t be used to compare absolute abundance levels between taxonomic groups.

| Fixed effects | Estimate | Std. Error | T-value | df <sup>2</sup> | P value |
| --- | --- | --- | --- | --- | --- |
| (Intercept) | -0.493 | 0.119 | -4.147 | 132.007 | <0.001 |
| landuse.L | -0.224 | 0.012 | -18.848 | 11624.206 | <0.001 |
| landuse.Q | -0.036 | 0.011 | -3.143 | 11648.378 | 0.002 |
| landuse.C | -0.025 | 0.011 | -2.381 | 11514.918 | 0.017 |
| taxon Plant | 0.380 | 0.190 | 1.999 | 135.141 | 0.048 |
| taxon Spider | 0.447 | 0.282 | 1.587 | 141.976 | 0.115 |
| records: taxon Bird | 0.074 | 0.009 | 8.013 | 7525.588 | <0.001 |
| records: taxon Plant | 0.095 | 0.014 | 7.010 | 8403.767 | <0.001 |
| records: taxon Spider | 0.318 | 0.033 | 9.664 | 8536.208 | <0.001 |
| landuse.L: records: taxon Bird | 0.173 | 0.010 | 17.445 | 12252.253 | <0.001 |
| landuse.Q: records: taxon Bird | 0.004 | 0.010 | 0.428 | 12989.143 | 0.668 |
| landuse.C: records: taxon Bird | 0.025 | 0.009 | 2.677 | 12037.901 | 0.007 |
| landuse.L: records: taxon Plant | 0.066 | 0.013 | 5.039 | 12260.474 | <0.001 |
| landuse.Q: records: taxon Plant | 0.017 | 0.012 | 1.454 | 11907.732 | 0.146 |
| landuse.C: records: taxon Plant | 0.012 | 0.011 | 1.085 | 12070.948 | 0.278 |
| landuse.L: records: taxon Spider | 0.041 | 0.034 | 1.221 | 15497.076 | 0.222 |
| landuse.Q: records: taxon Spider | -0.027 | 0.024 | -1.136 | 11451.153 | 0.256 |
| landuse.C: records: taxon Spider | -0.043 | 0.024 | -1.787 | 12218.887 | 0.074 |
| Random effects variance: residual = 0.245 , study = 1.043, species: study = 0.138 |  |  |  |  |  |
| Interclass correlation= 0.830 (i.e. proportion of total variance attributed to differences between groups) |  |  |  |  |  |
| Marginal R<sup>2</sup> = 0.022; Conditional R<sup>2</sup> = 0.832. |  |  |  |  |  |
| <sup>1</sup> log<sub>10</sub>(total abundance)~ landuse*(records:taxon)+ taxon+(1\|study/species) + offset(log<sub>10</sub>(total effort)) |  |  |  |  |  |
| <sup>2</sup> Satterthwaite approximation to degrees of freedom (Kuznetsova *et al.* 2017) |  |  |  |  |  |

### Extrapolation and aggregate indicators

PREDICTS studies have over-sampled the top end of the range of the number of records per species in GBIF as a whole (Fig. 1) — a feature also present in other leading biodiversity databases (fig. S3). Therefore, despite the uncertainty involved, it seems important to try to correct this undersampling. We corrected this by extrapolating the Records model to other named species in GBIF in the same taxonomic groups.

To obtain an average global measure of change in species abundance, we made predictions based on the main effects of the Records model for every named species in GBIF and every land-use type (even if that species-land use type combination did not appear in the raw data). Thus, we estimated the potential abundance for each of the 273,420 species for which number of records was available. This species vs land-use type matrix of predictions was used to calculate the geometric mean difference between primary vegetation and each other land-use type across species. We back-transformed this mean to give an approximate proportional change.

We placed confidence intervals on the predicted, proportional change by first, generating a large population of complete sets of plausible model parameters (N= 5000). We accounted for uncertainty in all the model fixed effects and their intercorrelations using the Cholesky decomposition, which produces random data that follow the covariance matrix of the Records model. Then, we calculated the predicted mean changes estimated with each of the 5000, parameter sets, and identified the 0.025 and 0.975 percentiles of the prediction distribution. The ‘confidence’ of this interval should be understood as the confidence in the average effect, not the confidence of what we would find sampling any one location or species.

### Robustness to data exclusion

As a supplementary analysis, we assessed the robustness of the ‘Records model’ to extrapolate abundance predictions. We set up a scenario within the PREDICTS dataset to exclude species with abundances closest to zero (assuming these species were most likely to have been missed if the sampling effort had been lower) and re-fit a ‘Training model’ to the remaining higher-abundance species. To do this, we first calculated each species’ mean abundance in each study. We then divided each study into a testing dataset, containing species with less than 10% of the study’s mean abundance (n=2,017 species-land use observations) and a training dataset, species with more than 10% of the mean abundance (n=16,681 observations). We used 10% as the threshold because this was approximately (assuming random resampling) the probability of a species being absent in at least two land uses, i.e. the proportion of zero abundance cases squared (0.322^2^= 0.10).

The parameter estimates of the Training model, i.e. model fitted with the training dataset (Table S6), fall within the 95% confidence interval of the estimates of the Records model (fitted with all observations; Fig. S3). The fixed effects of the Training model have the same pattern of significance as the Records model, except for the Plant taxon which is not significantly different from the Bird taxon. Finally, the correlation between the predicted and observed abundances in the testing dataset is similar for both models (Records model R^2^ =0.0017, Training model R^2^ =0.0019; Fig. S4). These results suggest that extrapolating to the low end of the recordability range is reasonable since the same trends seem to continue.

### Biodiversity intactness index comparison

We compared our estimations of change with those obtained through a leading global biodiversity indicator: the latest version of the Biodiversity Intactness Index (BII, Fig. 3b, Purvis 2019; De Palma *et al*. 2021; De Palma *et al*. 2024). We used the BII as a reference point for our prototype indicator, because it uses the PREDICTS database, and compares all land uses to primary vegetation as a reference level. If our prototype showed large differences from the BII, this would indicate a large impact of differing assumptions of the two models.

The BII combines a model of total abundance and a model of compositional similarity of species in a given area relative to primary vegetation (Newbold *et al*. 2016; De Palma *et al*. 2021). The total abundance is defined as abundance per unit effort. The composition similarity uses the balanced Bray-Curtis dissimilarity statistic– and this cannot be calculated for all the studies included in the total abundance statistic. The BII is obtained by multiplying the estimates of the two models for each land-use and intensity class, such that any shifts away from the composition of primary vegetation are negative, but these can be offset by increases in total abundance (for more details see De Palma *et al*. 2021; De Palma *et al*. 2024).

We used the land cover classification proposed by Outhwaite et al. and the species ‘Best guess binomial’ attribute to calculate compositional similarity. We obtained the BII index for the PREDICTS studies on birds, plants or spiders, as were used to fit our model (i.e. those with at least some Primary vegetation data and more than one species). We used 141 studies to estimate total abundance and 87 studies for compositional similarity (Table S7 and Table S8).

## Results

The ‘Records model’ shows that the local abundance of a species in PREDICTS is significantly affected by land use, the number of occurrence records in GBIF (hereafter ‘number of records’) and their interaction (Table 1 and Fig. 2). The baseline effect of land use (shown by the main effect terms in Table 1, and applicable to species with the average number of records) shows declining abundance in the order primary vegetation, secondary vegetation, low-intensity agriculture, high-intensity agriculture. The step down from low-intensity agriculture to high-intensity agriculture has the greatest magnitude (ΔPredicted log_10[_abundance] = −0.16; fig. S2).

**Figure 2.**
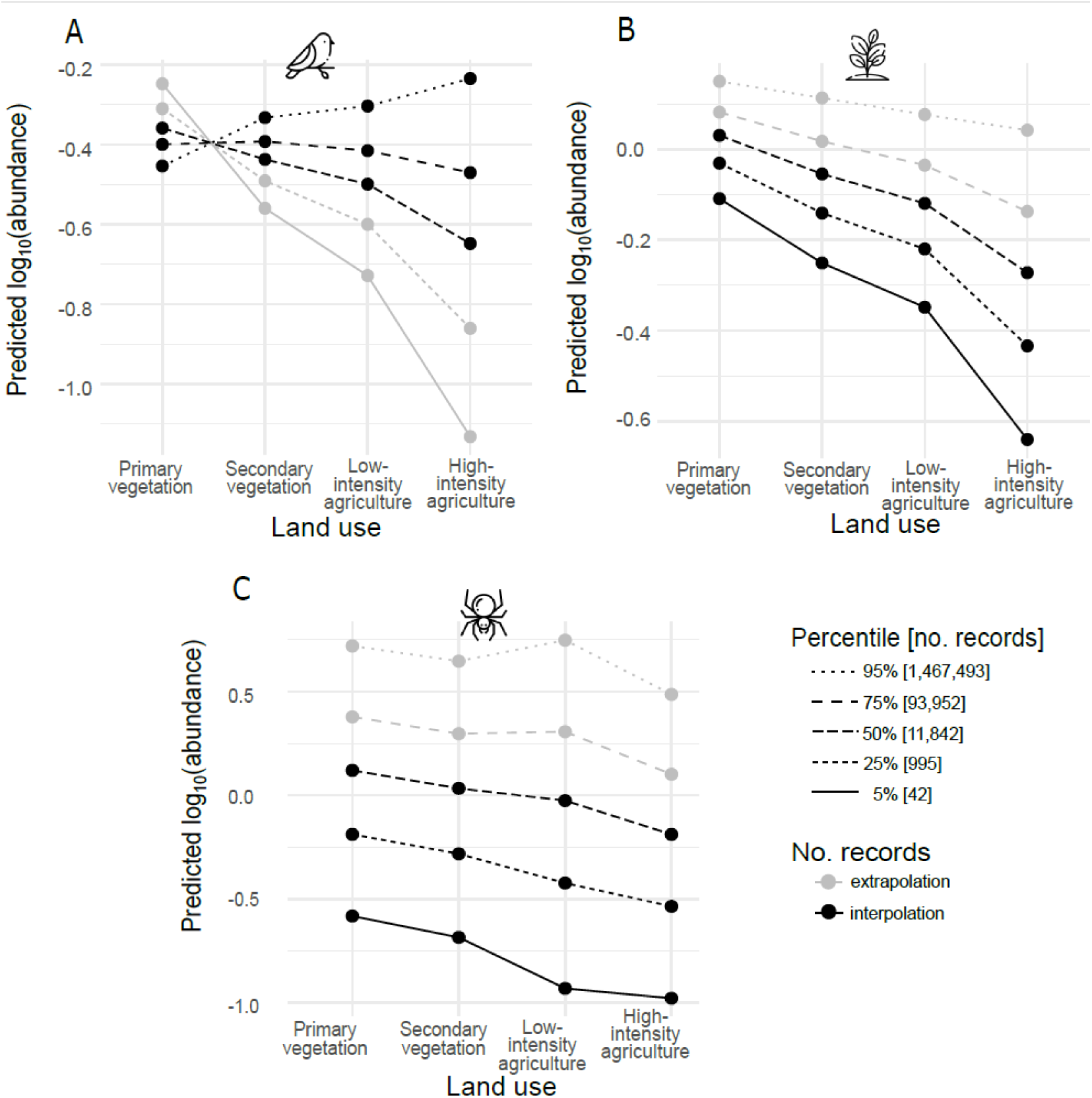
Predicted abundance by taxon, showing the interaction between land use and number of GBIF records for a species. Predictions are made for different percentiles of the distribution of the number of records for all observed (A) bird, (B) plant and (C) spider species (cf. grey density plot in Fig. 1a). In black (interpolation), predictions are within the range of numbers of records for the taxon; in grey (extrapolation), there are no species with this number of GBIF records within the taxon in our PREDICTS sample (table S4).ype or paste legend here. Paste figure above the legend.

The interaction between land use and number of records given taxonomic group is overall highly significant (ΔAIC= 399 compared to model without land use and records interaction, table S5), and the effect of these interactions is shown in Fig. 2. Overall, species with fewer occurrence records are consistently more negatively affected by anthropogenic land uses. For birds (Fig. 2a) and plants (Fig. 2b), as the number of records for a species increases, the relationship with land use becomes shallower (more positive) and more linear (quadratic and cubic terms closer to zero). For spiders, the interaction terms are individually not significant, but the trend is qualitatively similar as for the other groups. For example, like in bird and plant species, spider species (Fig. 2c) at the lowest percentile number of records show the greatest total decline between primary vegetation and intensive agriculture. Only in birds with among the highest observed numbers of records do we see a progressive increase in abundance across the land use categories, representing species that benefit from anthropogenic factors (Fig.2A). For plants and spiders, the highest prediction lines are extrapolation outside the range of numbers of records observed in GBIF for these taxa (grey lines in Fig2B-C), and we do not predict a positive response to anthropogenic land use anywhere in our parameter space.

We converted our results to an overall summary of relative abundance of organisms in modified land uses compared to primary vegetation (Fig. 3a, note the un-logged y-axis in comparison to Fig. 2): specifically, this took the geometric mean across species of their model-predicted abundances in each land use type.

**Figure 3.**
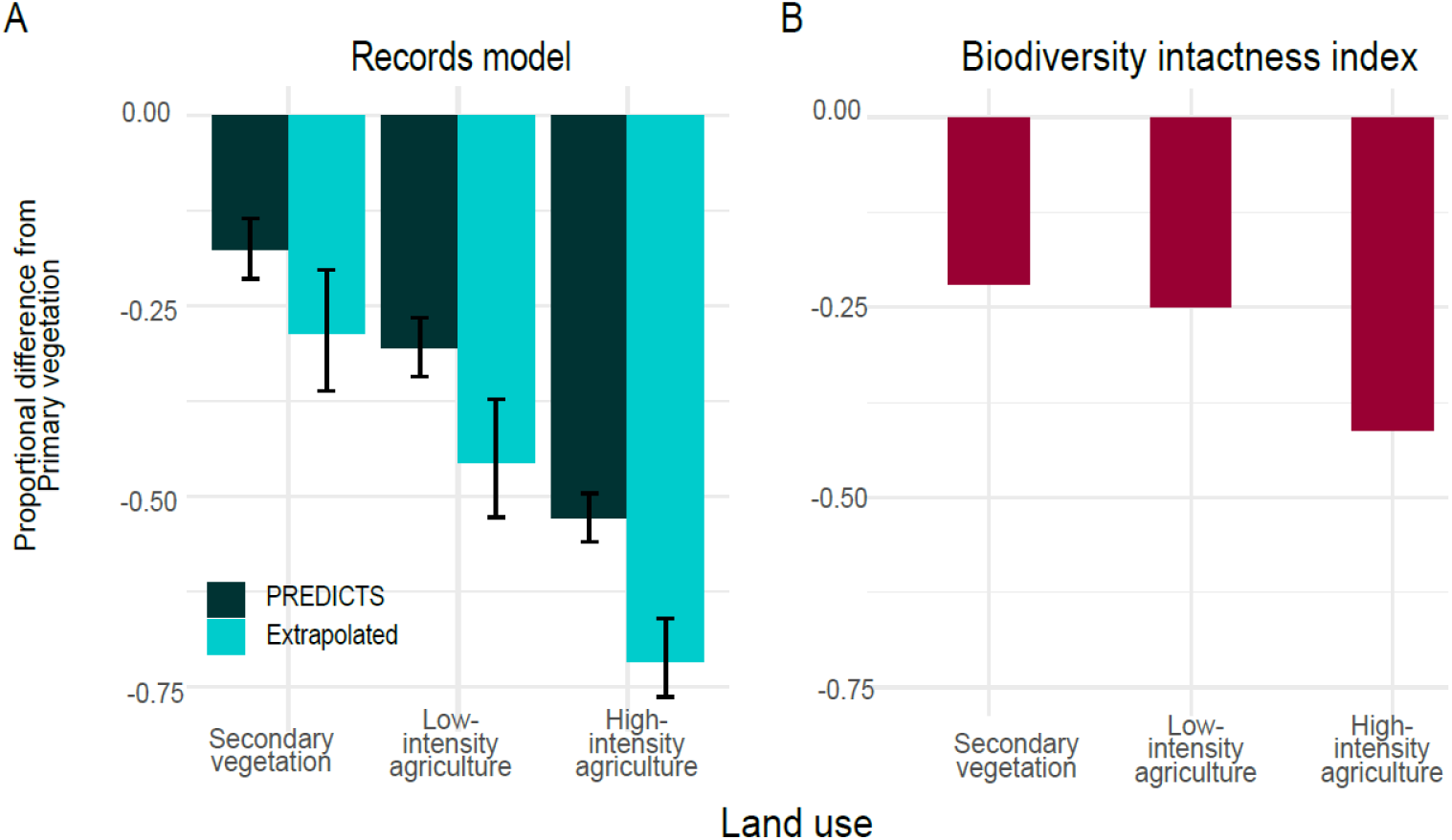
Proportional difference of three land cover types to primary vegetation. (A) Difference of the geometric mean predicted abundance of the Records model (95% CI), across the 4,454 bird, plant and spider species observed in the PREDICTS database (dark green) compared to the difference estimated by extrapolating to all 273,420 species for which the number of records was available in GBIF (light green). (B) The Biodiversity Intactness Index (methods in De Palma *et al*. 2024) based on PREDICTS database studies of birds, plants, and spiders.

For all modified land uses, the proportional declines estimated after extrapolating the Records model to other named species in GBIF were considerably greater than those estimated using only the species included in the PREDICTS database (Fig. 3a). The percentage-point difference made by extrapolation was 11% for secondary vegetation, 15% for low-intensity agriculture and 19% for high-intensity agriculture, which, for example, makes high-intensity agriculture’s predicted proportional decline 71.6% (95% CI, 66 to 76).

We then compared the mean proportional decline estimates of the Records model to the ones obtained through the Biodiversity Intactness Index (BII). Although ecologists will readily perceive differences between these metrics for comparing land uses, when presented to a policymaker, they could be used to answer the same question: “How bad is it?”. The fairest way to compare our metric to the BII for its optimism/pessimism is to use the Records model restricted to PREDICTS observations (Fig.3A ‘Records model, Predicts’) vs. the estimates derived from the BII model for the same observation set (Fig. 3B). This comparison shows the BII is more optimistic about the change to both agricultural land uses but slightly more pessimistic about the change from primary to secondary vegetation (Fig. 3).

## Discussion

Hard-to-sample species are seriously under-represented in the PREDICTS database, and some other databases (fig. S3) that are otherwise extremely useful to document biodiversity loss and attribute its causes. We found that — consistently across taxonomic groups — hard-to-sample species are more negatively affected by anthropogenic land uses. No global metrics are immune from these biases (e.g. those reviewed in Rosa *et al*. 2020). Therefore, in the future development of all these metrics, the feasibility of extrapolating to under-sampled and potentially more sensitive species should be seriously considered.

The generalisable message of our study transcends the specific decline metric that we chose and its comparison to the BII. However, it is reassuring that the trends of our metric are broadly similar to the BII when calculated on similar data: i.e. it provides a ‘sense check’ of our metric. If we could find a way to extrapolate the BII to account for under-recorded species, we would expect the values to decrease, because of the stark bias shown in Fig. 1. We have not attempted any extrapolation method for the BII here, since it would require some complex additional assumptions. In brief, the statistical models underlying the BII do not contain any species-specific values that can be straightforwardly extrapolated to an additional set of species (for details on the BII’s methods see De Palma *et al*. 2024).

We have shown that statistical extrapolation is viable, using information from GBIF, and the difference it makes seems big enough to matter to policymakers (over 10 points on a 0-100 scale). We argue that an extrapolation method is the most powerful way to address the under-sampling of species. Such a method does not have to be the exact one that we chose.

Despite their limitations, both PREDICTS and GBIF contain vital information for understanding biodiversity loss, and prioritising action towards nature’s recovery. We are not here simply adding to the list of limitations of such data collection, which have been extensively discussed elsewhere (Troia & McManamay 2016; Hughes *et al*. 2021). We are showing how, when used together, the strengths of each database can complement the other. The main strength of PREDICTS is its structured design, where equivalent surveys were applied in different land uses and/or intensities (Newbold *et al*. 2019; Purvis 2019). However, the studies within PREDICTS mostly use community sampling methods with a level of effort that could not be expected to sample all the rarest species in the habitat concerned. Though PREDICTS should continue to grow, it will still consist of patchy snapshots of communities (Hudson *et al*. 2014). Standardised field surveys are labour-intensive and technically demanding (Gotelli *et al*. 2023), so the amount of survey effort required to extend PREDICTS studies to capture rarer species is unfeasible. By contrast, extrapolating from these studies is highly feasible as we have shown.

The main strength of GBIF is its sheer size and taxonomic coverage. However, it is a collection of heterogeneous occurrence data sets, collected for different reasons, and species that are rare in different ways can sometimes be under- and sometimes over-represented (Troia & McManamay 2016; Hughes *et al*. 2021; Garcia-Rosello *et al*. 2023). For instance, of the species included in our analyses, the spider taxon is estimated to contain four times as many un-named species (Agnarsson *et al*. 2013), whereas the taxonomy of higher plants and birds is relatively comprehensive (Hobohm *et al*. 2019; Clements 2023). If we had been able to include species entirely missing from GBIF, our extrapolated estimates of abundance change would have been even more pessimistic. However, these estimates of the numbers of missing species are themselves extrapolations, and the compounded uncertainty would be very high.

We propose that the number of records for species in GBIF is a useful metric of their ‘recordability’. If recordability just means a species’ frequency of recording in other biodiversity databases, our investigations support this strongly. However, two questions deserve further investigation: first, is recordability a consistent feature of a species? If not, it may be necessary to decompose recordability into different contributing factors such as range size (fig. S6), local abundance, and the probability of detection under commonly used ecological survey methods. Second, if it is a consistent feature, are there better ways to measure it than counting GBIF records? For example, note that we have fitted relationships independently for each taxon, because taxonomic group impacts almost everything about how occurrence records are collected, and by whom. If we could find a metric of recordability that accounted for group-wise differences in recording culture, we might find this more reliably related to the response to land-use, and therefore more robust to extrapolate with.

If we wish to extrapolate to the greatest possible number of hard-to-sample species, we have little choice but to use GBIF data. Because of GBIF’s integration with the Catalogue of Life (the global species checklists aimed to include all known species of organisms on Earth), there is no better source for a complete list of accepted species for most taxonomic groups. We may believe that functional, ecological traits underlie species’ responses to land use, and these traits happen to be correlated to recordability, but trait databases are only developed for well-studied taxa (Table S9, Sandel *et al*. 2015; Etard *et al*. 2020). For example, Sykes et al. (Sykes *et al*. 2020) found land-use responses could be related to three aspects of rarity among vertebrate species, but in fact, around three-quarters of the species included were birds, and the third aspect of rarity could only be determined for about a quarter of the species. Thus, while such studies are valuable for exploring ecological mechanisms, they are currently less useful for developing representative global indicators. Note that the obvious trait of range size (fig. S6, table S9) could be estimated from GBIF data, but these estimates themselves would be less reliable for under-recorded species. The urgent need for conservation decisions cannot wait for the ideal data to be available (Garcia-Rosello *et al*. 2023), so we need to maximize the utility of the data we currently have.

There are two alternative ways forward for improving global indicators to better account for hard-to-sample species. One would be to further develop our indicator shown in Fig. 3a, and the other to develop extrapolation methods for the other established indicators. To the extent that other indicators have already built up political support, it may be more fruitful to adapt them. There is already a history of adapting indicators where better data and fitting methods become feasible (Ledger *et al*. 2023); and although there are good reasons for measuring some differing aspects of biodiversity, policymakers can get frustrated if indicators seem inconsistent (Hill *et al*. 2016). Extrapolation may even make a difference to the indicators that already restrict themselves to vertebrates and plants (e.g. several of the indicators used for intercomparison in Pereira *et al*. 2024) if they don’t already model all species in the group. However, we would argue strongly for the inclusion of a greater range of taxa in any indicators where possible; for example, the BII is particularly well designed for including many, diverse taxa (De Palma *et al*. 2021).

Our prototype indicator could be extended in several ways where there is enough data to support this, for example including more taxonomic groups, more land-use categories, or having variants for different continents or biomes. We could also relax our assumption about weighting every species equally in the geometric mean. Before making any refinements, however, it is important to consider how the indicator would be used, and interpreted by policymakers (Stevenson *et al*. 2021). Some indicators are great for raising awareness and convincing non-scientists of the need for action (Ledger *et al*. 2023). However, the best indicators for deciding between specific policy options are not necessarily the same (Nicholson *et al*. 2012; Hill *et al*. 2016). Although there are calls for causal models that could work equally well for communication, policy testing, monitoring and evaluation (Nicholson *et al*. 2012; Hill *et al*. 2016; Gonzalez *et al*. 2023), there are still many hurdles to overcome for this to happen globally. Our approach to extrapolation is quite general, so it could be applied as part of a causal model to help correct for patchy data availability. It could be applied to develop a global model with as many species as possible, but simple land cover categories because these are the only ones that are mapped globally (Hill *et al*. 2018). Or it could be used for national/regional decision-making with only the species that occur locally and made more relevant to the land-use transitions that can occur locally (Martin *et al*. 2022).

The fact that a large fraction of the world’s species are under-recorded is often mentioned, but rarely do studies suggest practical workarounds for correcting for the biased assessments that may arise from biased data —they may more often suggest collecting more data (Gonzalez *et al*. 2023). Here, we present a workaround based on statistical extrapolation and show the magnitude of difference this could make across selected taxonomic groups. Based on the strong sampling biases we have uncovered against hard-to-record species, we suggest that all developers of high-level biodiversity indicators attempt to implement an extrapolation solution that is relevant to their outcome of interest.

## Acknowledgements

We thank Samantha Hill for her advice on analysing PREDICTS data, Simon Smart for his feedback in the early stages of the project, and Michael Begon and Jane Rees for their suggestions to improve the manuscript.

