## Supplementary Information for "Hard-to-sample species are more sensitive to land-use change: implications for global biodiversity metrics"

**This PDF file includes:**

Supporting text
Figs. S1 to S7
Tables S1 to S9
SI References

**Exploratory analyses**

We performed an initial exploration on the availability of data potentially related to species' sensitivity to land use change (Table S9). We obtained the IUCN Red List estimations (5) of the Extent of Occurrence (EOO; area contained within the shortest continuous imaginary boundary which can be drawn to encompass all the known, inferred or projected sites of present occurrence of a taxon) and Area of Occupancy (AOO; area of suitable habitat currently occupied by the taxon). We found EOO data for only 13% of the bird species and less than 1% of plants and spiders in our GBIF species sample (Table S9), and less than 1% of AOO data for all groups. Nevertheless, we found a positive strong correlation between EOO and number of records (Figure S6).

Additionally, we compared the BII estimations of change when calculated with our studies' sample (bird, plant and spider studies) with those obtained when including all suitable studies to calculate the BII for all taxonomic groups. For the latter, we modelled total abundance with 79% of the observations in PREDICTS (720 studies), and compositional similarity with 39% of the observations (294 studies; Table S7 and Table S8). The estimates of loss to secondary vegetation for all taxonomic groups (21%) and bird, plant and spider species (22%) are very similar (Fig. S7). However, these estimates vary considerably for the other land uses. The difference estimated between primary vegetation and agricultural land uses is similar regardless of use intensity, when considering all taxonomic groups (low-intensity agriculture 41%; high-intensity agriculture, 43%). This contrasts with the estimations obtained for bird, plant and spider studies, where the decrease for low-intensity agriculture (25%) is considerably lower to that of high-intensity agriculture (41%).

**Fig. S1**

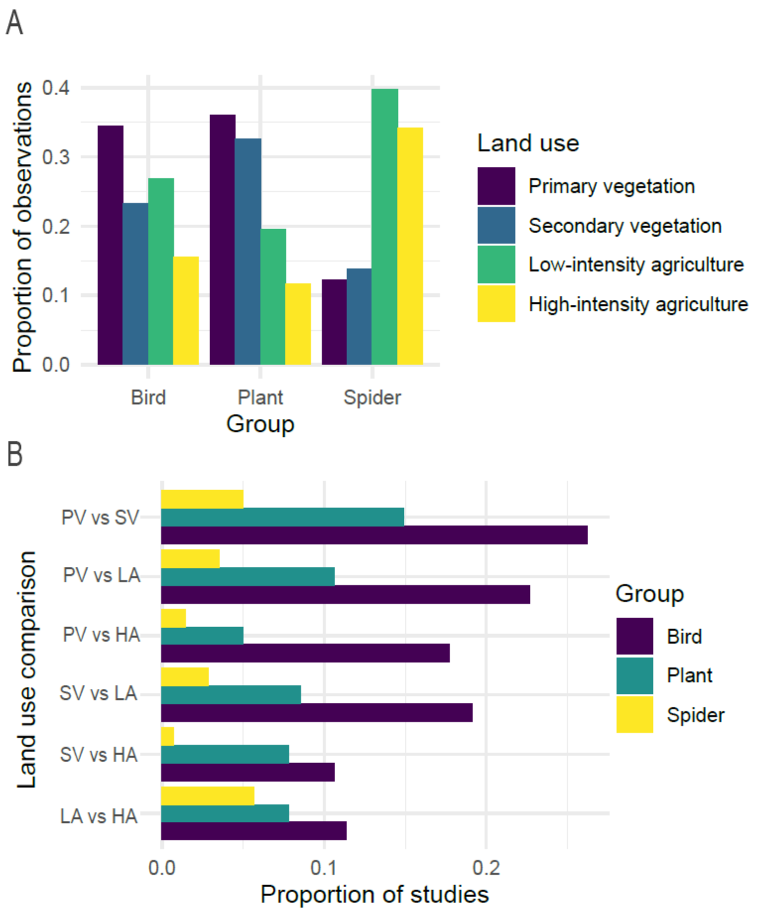


**Fig. S1.** The balance of observations and studies available for different taxa and land uses in the PREDICTs subset used to fit the Records model. (A) Proportion of the taxon's total of observations in each land use type. Note that the total observation number differs for each taxon: birds (n= 11,822; 63%), plants (5,362; 29%) and spiders (1,514; 8%). (B) Proportion of the grand total of studies (n=141) by pair-wise land use comparison and taxon (totals to more than 1 due to studies that made more than one comparison). PV: Primary vegetation; SV: Secondary vegetation; LA: Low-intensity agriculture; HA: High-intensity agriculture.

**Fig. S2**

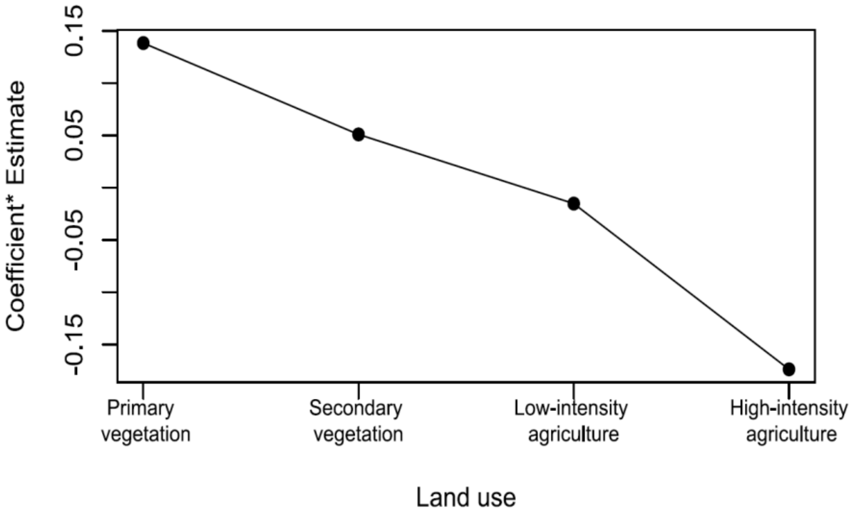


**Fig. S2.** Land-use change baseline effect on abundance. Effect estimated from the linear (landuse.L), quadratic (landuse.Q), and cubic (landuse.C) estimates for the main effect in the Records model (i.e. rows 2-4 in Table 1). This can be understood as the average effect over the three taxonomic groups if they have the mean log number of GBIF records.

**Fig. S3**

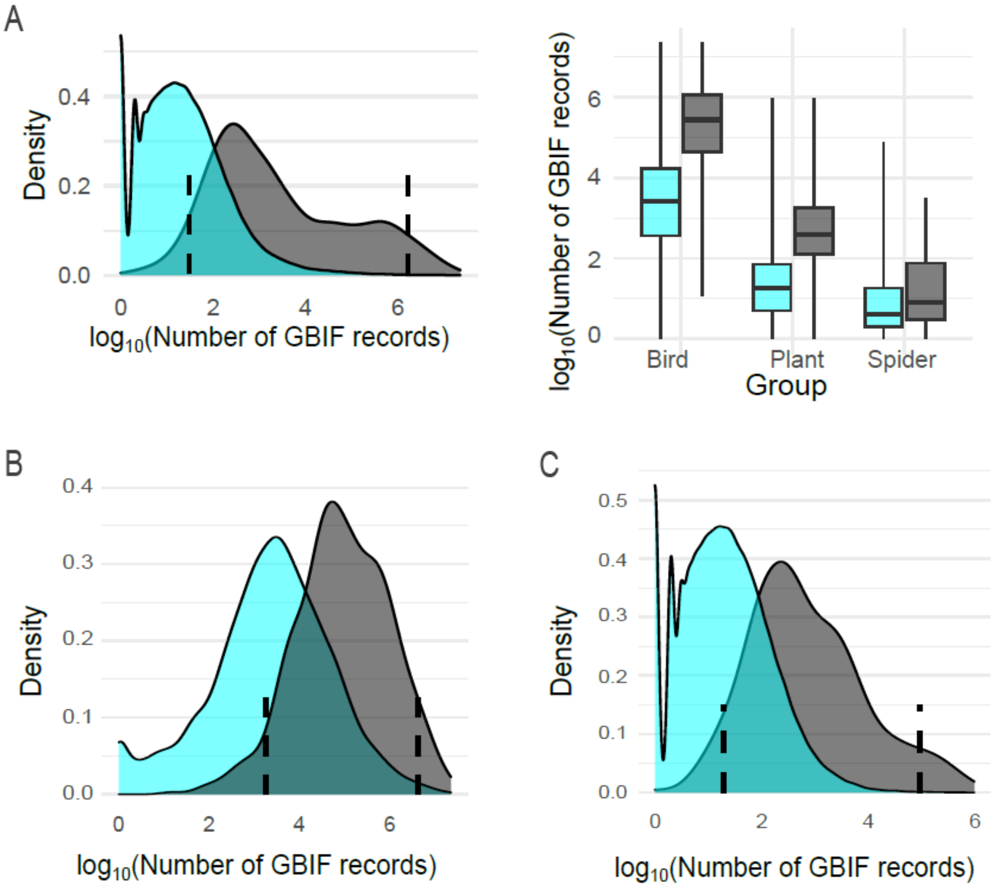


**Fig.** S3. Distribution of number of GBIF records. Distributions as a density plot comparing bird, plant and spider species available in GBIF (cyan) to the species also present in other leading biodiversity databases (grey). (A) Left: GBIF (n=273,420) vs BioTIME (n=2,448), right: distributions summarised as boxplots for each taxonomic group (1); (B) GBIF (n=10,334) vs Living Planet(n= 1,072), birds only (2); (C) GBIF (n= 262,526) vs COMPADRE(n=329), plants only (3). Dashed lines indicate the 0.05 and 0.95 percentiles of the distributions in grey.

**Fig. S4**

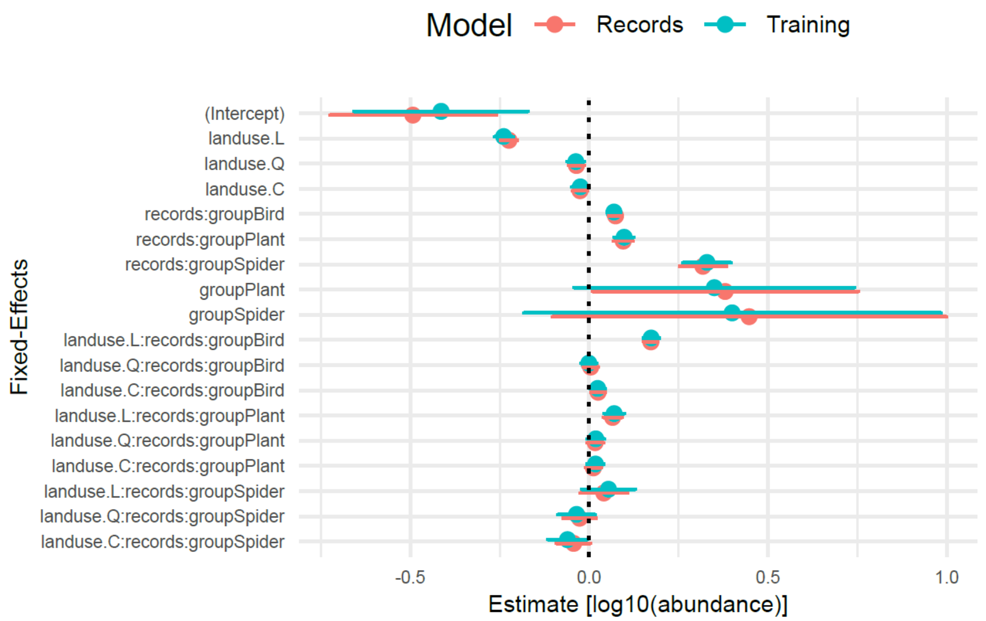


**Fig. S4.** Fixed effects estimate with or without a subset of low-abundance species**.** Comparison between the estimates and respective 95% confidence intervals of the fixed effects of the 'Records model', including low-abundance species, and 'Training model', excluding low abundance species.

**Fig. S5**

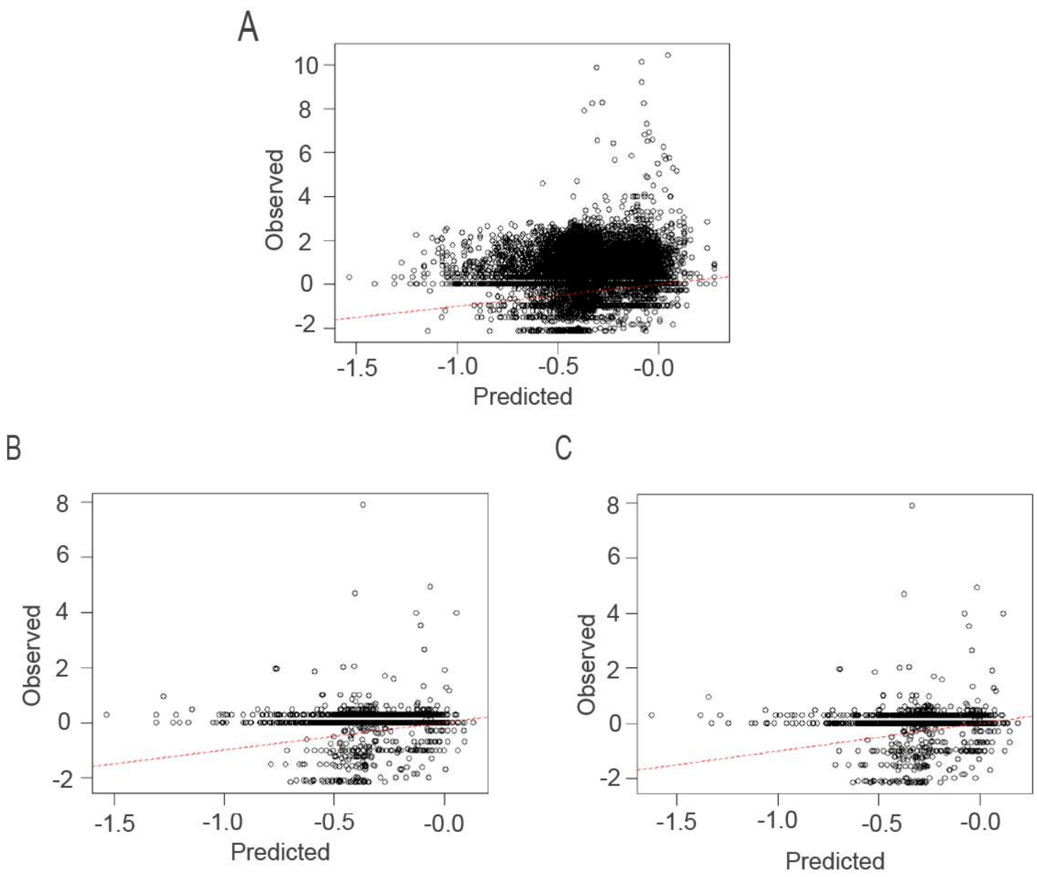


**Fig. 5.** Testing extrapolation in terms of observed vs predicted abundance**.** (A) Predicted values using fixed effects of the Records models vs observed values of all data used in fitting it (n= 18,698; $R^2$=0.0093). (B) Predicted values using Records model vs observed values of testing data, i.e. observation with <10% mean abundance per species by study (n=2,017; $R^2$= 0.0017). (C) Predicted values using the Training model (fitted with > 10% mean abundance per species by study, n= 16,681) vs observed values of testing data (n=2,017; $R^2$= 0.0019). Dotted line indicates 1:1 relationship, i.e. expectation if observed values were identical to predicted values.

**Fig. S6**

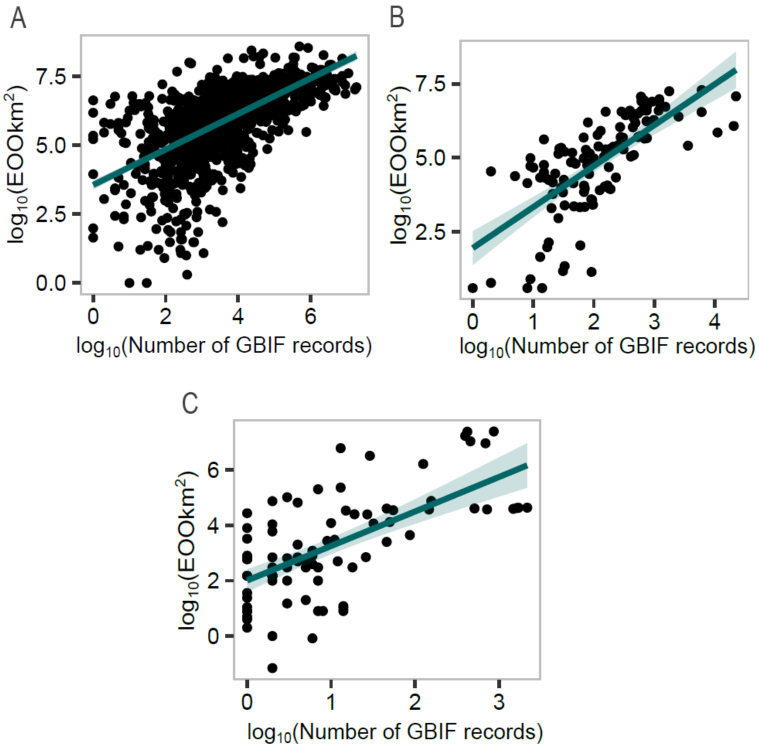


**Fig. S6.** Relationship between the number of records (GBIF occurrence count) and the IUCN Red List estimations of the Extent of Occurrence (EOO; 4)**.** Solid line = linear model regression, shaded region = 95% confidence interval. (A) Birds, estimate= 0.64, p< 2.2e-16; (B) plants, estimate = 1.38, *p*< 2.2e-16. See Table S9 for the proportion of species for which this comparison is possible; (C) spiders, estimate= 1.25, p< 5.9e-12.

**Fig. S7**

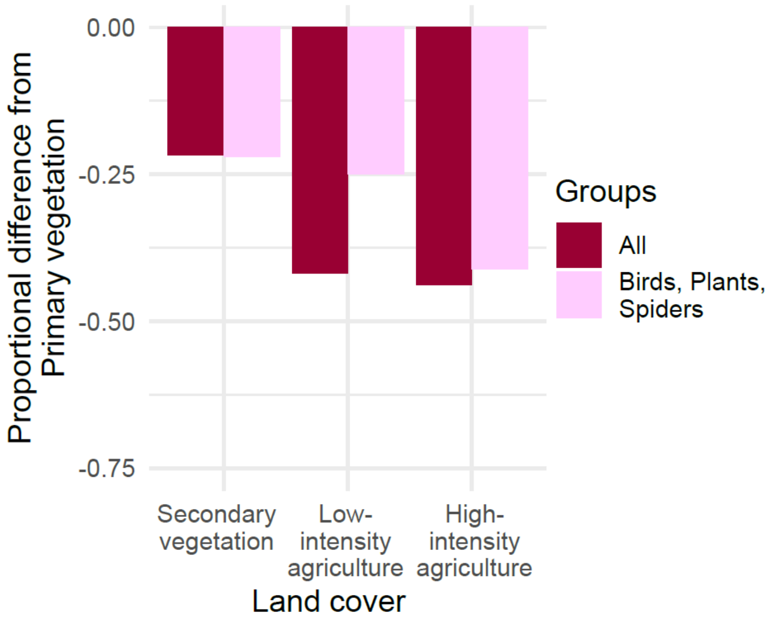


**Fig. S7.** Proportional difference of three land cover types to primary vegetation**.** Differences estimated with the Biodiversity intactness Index including relevant PREDICTS' studies (Table S8) for all taxonomic groups (dark red), or studies of birds, plants, and spiders (pink); for details on BII's methods see (5).

**Table S1**

**Table S1.** Land use classes. PREDICTS database land use and use intensity classes (6) and simplified classification suggested by Outhwaite, McCann and Newbold (7).

| PREDICTS | Simplified classification |
|---|---|
| Primary vegetation —minimal, light, and intense use | Primary vegetation |
| Secondary vegetation (mature, intermediate, young, and indeterminate age) —minimal, light, and intense use | Secondary vegetation |
| Plantation forest — minimal use<br>Pasture — minimal use and light use<br>Cropland — minimal use and light use | Low-Intensity agriculture |
| Plantation forest — light and intense use<br>Pasture — intense use<br>Cropland — intense use | High-Intensity agriculture |
| Urban<br>Cannot decide | Excluded |

**Table S2**

**Table S2.** Metrics of dataset size. Number of studies in PREDICTS that report an abundance metric, the taxa of interest were identified at the species level and compare two or more of the land uses; number of 'total abundance' observations (species times land uses by study) and species reported in these studies. For bird, plant and spider studies (Subset studies), number of species for which it was possible to obtain the number of records from GBIF and the number of species in PREDICTS whose name matches GBIF species.

| Taxon | Studies | Observations | Species | Total Spp. with records in GBIF in the taxon | Spp. in PREDICTS with a name that matches to GBIF |
|---|---|---|---|---|---|
| All studies | 413 | 67,000 | 17,986 | - | - |
| Subset studies | 141 | 35,637 | 9,212 | - | 4,454 |
| Birds | 75 | 16,763 | 3,100 | 9,381 | 2,188 |
| Plants | 49 | 16,289 | 5,510 | 242,269 | 1,916 |
| Spiders | 17 | 2,585 | 602 | 21,770 | 350 |

**Table S3**

**Table** S3. Percentiles of the number of records. Percentiles of the number of records for the three and each of the taxa (birds 63%, plants 29%, spiders 8%) in the PREDICTS' sample, in square brackets the transformed number of records = [$\log_{10}$ (number of GBIF records) - mean ($\log_{10}$(number of GBIF records))], where mean = 3.979.

| | 25% | 50% | 75% | 95% |
|---|---|---|---|---|
| Three taxa | 995<br>[ -0.981] | 11,842<br>[0.094] | 93,952<br>[0.994] | 1,467,493<br>[2.188] |
| Birds | 11,184<br>[0.069] | 52,525<br>[0.741] | 211,424<br>[1.346] | 2,734,213<br>[2.458] |
| Plants | 89<br>[-2.029] | 314<br>[-1.481] | 1,236<br>[-0.887] | 28,780<br>[0.480] |
| Spiders | 645<br>[-1.169] | 3,105<br>[-0.487] | 6,068<br>[-0.196] | 20,801<br>[0.339] |

**Table S4**

**Table S4.** Number of pair-wise comparisons of land cover types. Number of studies within the 141 subset PREDICTS studies that could be analysed along with GBIF record number for bird plant or spider species. See also Fig. S1

| | Primary Vegetation | Secondary vegetation | Low intensity agriculture |
|---|---|---|---|
| Secondary vegetation | 65 | - | - |
| Low-intensity agriculture | 52 | 43 | - |
| High-intensity agriculture | 34 | 27 | 35 |

**Table S5**

**Table S5.** Comparing the Records model to simpler variants with fewer fixed effects. Terms considered were land use type (landuse), transformed number of species records (records) and taxonomic group (taxon). Variance of random effects includes study and species within study (1|study/species).

| Fixed effects included | AIC | ΔAIC | Random effects variance | |
|---|---|---|---|---|
| landuse * (records : taxon) + taxon | 34100 | 0 | species: study<br>study<br>Residual | 0.138<br>1.043<br>0.245 |
| landuse * taxon + (records : taxon) | 34412 | 312 | species: study<br>study<br>Residual | 0.135<br>1.067<br>0.253 |
| landuse + taxon + (records : taxon) | 34499 | 399 | species: study<br>study<br>Residual | 0.134<br>1.054<br>0.255 |
| landuse * taxon | 34576 | 476 | species: study<br>study<br>Residual | 0.140<br>1.046<br>0.253 |

**Table S6**

**Table S6.** Summary of ‘Training model’[1]: structured the same as the Records model (Table 1), except that some low-abundance species have been excluded from the data. ‘landuse’ is an ordered factor, fitted with polynomial contrasts. ‘taxon’ is an unordered factor and the transformed number of GBIF records ‘records’ is a scalar. Number of observations = 16,681; studies = 137, species within studies=6,923.

| Fixed effects | Estimate | Std. Error | T-value | df[2] | P value |
|---|---|---|---|---|---|
| (Intercept) | -0.414 | 0.124 | -3.333 | 127.137 | 0.001 |
| landuse.L | -0.239 | 0.013 | -18.355 | 10502.875 | <0.001 |
| landuse.Q | -0.037 | 0.012 | -3.006 | 10530.820 | 0.003 |
| landuse.C | -0.025 | 0.012 | -2.107 | 10395.490 | 0.035 |
| taxon Plant | 0.350 | 0.200 | 1.747 | 129.814 | 0.083 |
| taxon Spider | 0.400 | 0.299 | 1.339 | 134.011 | 0.183 |
| records: taxon Bird | 0.069 | 0.009 | 7.331 | 6703.635 | <0.001 |
| records: taxon Plant | 0.098 | 0.014 | 7.048 | 7562.513 | <0.001 |
| records: taxon Spider | 0.329 | 0.034 | 9.611 | 7912.833 | <0.001 |
| landuse.L: records: taxon Bird | 0.173 | 0.011 | 16.147 | 11120.675 | <0.001 |
| landuse.Q: records: taxon Bird | -0.001 | 0.011 | -0.086 | 11829.972 | 0.932 |
| landuse.C: records: taxon Bird | 0.024 | 0.010 | 2.334 | 10907.529 | 0.020 |
| landuse.L: records: taxon Plant | 0.070 | 0.014 | 4.940 | 11142.076 | <0.001 |
| landuse.Q: records: taxon Plant | 0.018 | 0.013 | 1.472 | 10812.924 | 0.141 |
| landuse.C: records: taxon Plant | 0.018 | 0.012 | 1.477 | 10937.927 | 0.140 |
| landuse.L: records: taxon Spider | 0.054 | 0.038 | 1.411 | 14693.714 | 0.158 |
| landuse.Q: records: taxon Spider | -0.035 | 0.026 | -1.329 | 10258.768 | 0.184 |
| landuse.C: records: taxon Spider | -0.061 | 0.028 | -2.178 | 11354.143 | 0.029 |
| Random effects variance: residual = 0.261 , study =1.127, species: study = 0.115<br>Interclass correlation = 0.830; Marginal $R^2$ = 0.020; Conditional $R^2$ = 0.830 | | | | | |
| [1] log10(total abundance)~ landuse*(records: taxon)+ taxon+(1\|study/species) + offset(log10(total effort))<br>[2] Satterthwaite approximation to degrees of freedom | | | | | |

**Table S7**

Table S7. Species, observations, and studies representation in the PREDICTS database used to calculate the Biodiversity Intactness Index for the simplified landuse categories in Table S1.

| Used for the abundance sub-model[1] | | | |
|---|---|---|---|
| | Species | Observations | Studies |
| All studies | 29,221 | 3,658,623 | 720 |
| Subset studies | 9,802 | 1,125,195 | 141 |
| Birds | 3,145 | 481,423 | 75 |
| Plants | 6,042 | 559,220 | 49 |
| Spiders | 615 | 84,552 | 17 |
| Used for the compositional similarity sub-model[2] | | | |
| All studies | 16,279 | 1,659,453 | 294 |
| Subset studies | 7,601 | 824,588 | 87 |
| Birds | 2,477 | 348,150 | 46 |
| Plants | 4,842 | 444,704 | 32 |
| Spiders | 284 | 31,734 | 9 |

[1]Studies where the diversity metric was 'Abundance' and have sites in land covers of interest.
[2]Studies with some primary vegetation information and more than one species

**Table S8**

**Table S8.** Number of studies with a pair-wise comparison of land cover types. Studies in all PREDICTS database used to calculate the Biodiversity Intactness Index. In brackets, number of bird, plant and spider studies with a pair-wise comparison. Can be compared with Table S4.

| | Primary vegetation | Secondary vegetation | Low intensity agriculture |
|---|---|---|---|
| Used for the abundance sub-model | | | |
| Secondary vegetation | 222 [65] | - | - |
| Low-intensity agriculture | 148 [52] | 157 [43] | - |
| High-intensity agriculture | 104 [34] | 99 [27] | 125 [35] |
| Used for the compositional similarity sub-model | | | |
| Secondary vegetation | 160 [51] | - | - |
| Low-intensity agriculture | 109 [46] | - | - |
| High-intensity agriculture | 72 [29] | - | - |

**Table S9**

**Table S9.** Comparison of data availability for relevant traits to assess species sensitivity to land use change, by taxonomic group. The number of species with number of records in GBIF (GBIF); the number of these species with information on the extent of occurrence (EOO); area of occupancy (AOO; 4); and species with EOO estimates also present the PREDICTS database.

| Taxon | GBIF | EOO [%] | AOO [%] | EOO & PREDICTS [%] |
|---|---|---|---|---|
| Birds | 10,334 | 1,239 [12.0] | 76[0.74] | 369 [3.57] |
| Spiders | 29,117 | 159 [0.55] | 153[0.53] | 13 [0.04] |
| Plants | 262,526 | 127 [0.05] | 94 [0.04] | 7 [0.003] |
